# Pre- and post-task resting-state differs in clinical populations

**DOI:** 10.1101/2022.09.20.508750

**Authors:** Cindy Sumaly Lor, Mengfan Zhang, Alexander Karner, David Steyrl, Ronald Sladky, Frank Scharnowski, Amelie Haugg

## Abstract

Resting-state functional connectivity has generated great hopes as a potential brain biomarker for improving prevention, diagnosis, and treatment in psychiatry. This neuroimaging protocol can routinely be performed by patients and does not depend on the specificities of a task. Thus, it seems ideal for big data approaches that require aggregating data across multiple studies and sites. However, technical variability, diverging data analysis approaches, and differences in data acquisition protocols might introduce heterogeneity to the aggregated data. Besides these technical aspects, the psychological state of participants might also contribute to heterogeneity. In healthy participants, studies have shown that behavioral tasks can influence resting-state measures, but such effects have not yet been reported in clinical populations. Here, we fill this knowledge gap by comparing resting-state functional connectivity before and after clinically relevant tasks in two clinical conditions, namely substance use disorders and phobias. The tasks consisted of viewing craving-inducing and spider anxiety provoking pictures that are frequently used in cue-reactivity studies and exposure therapy. We found distinct pre- vs. post-task resting-state connectivity differences in each group, as well as decreased thalamo-cortical and increased intra-thalamic connectivity which might be associated with decreased vigilance in both groups. Notably, the pre- vs. post-task thalamus-amygdala connectivity change within a patient cohort seems more pronounced than the difference of that connection between the smoker vs. phobia clinical trait. Our results confirm that resting-state measures can be strongly influenced by changes in psychological states that need to be taken into account when pooling resting-state scans for clinical biomarker detection. This demands that resting-state datasets should include a complete description of the experimental design, especially when a task preceded data collection.

## 1. Introduction

Resting-state functional brain connectivity is a promising tool for the development of potential biomarkers for improving prevention, diagnosis, and treatment in psychiatry. It is a task-free neuroimaging protocol that patients can generally perform easily irrespective of the type or the severity of their condition. Indeed, in recent years, candidate resting-state biomarkers have been reported in nearly all psychiatric conditions. For example, based on resting-state data, relatively high levels (> 80%) of classification accuracy have been achieved for psychiatric conditions that include depression (Drysdale et al., 2017), post-traumatic-stress disorder (Nicholson et al., 2019, 2020), consciousness states (Campbell et al., 2020), or tobacco use disorder (Wetherill et al., 2019).

Another key advantage of resting-state biomarkers is that they do not depend on the specificities of a task. Consequently, they appear ideal for between-study comparisons and for collapsing datasets from multiple studies and multiple sites, which would improve generalizability and statistical power. This is of critical importance, as correlation-based approaches are gradually replaced by predictive machine-learning methods, which, although more powerful, require large amounts of data (Khosla et al., 2019). Pooling multiple resting-state studies is increasingly popular (Abraham et al., 2017; Tanaka et al., 2021; Turner, 2013), but it is far from trivial, as shown by the modest success of recent multisite studies. Classification algorithms of major depressive disorder (Xia et al., 2019) and schizophrenia (Cai et al., 2020), for instance, poorly generalized from one study to another despite high classification accuracy within single studies.

Importantly, one should not take for granted that the gain from increasing sample size will systematically outweigh the introduction of heterogeneity (Bari et al., 2019). Heterogeneity can be due to technical variability (e.g. scanner type, field strength, imaging sequences…) (Yamashita et al., 2019), different data analysis approaches such as functional connectivity, amplitude of low frequency fluctuations, graph theory, clustering algorithms and pattern classification, as well as differences in data acquisition protocols. Despite being task-free, resting-state protocol variables such as acquisition duration and whether or not the participants should fixate or close their eyes, were intensely researched and discussed (Agcaoglu et al., 2019). However, heterogeneity can also be due to differences in transient psychological states. Although many studies have tried to predict rather stable traits, resting-state measures may not display the same level of within-subject stability. Indeed, several studies on healthy participants reported resting-state functional connectivity changes to the hippocampus after an associative memory task (Tambini et al., 2010), to the dorsal attention network, default mode network, and visual networks after a perceptual learning task (Lewis et al., 2009), to motor areas after a finger-tapping task (Sarabi et al., 2018), or to the olfactory piriform cortex after an olfactory task (Cecchetto et al., 2019; Sarabi et al., 2018)). This indicates that the psychological state of the participants alters resting-states. A similar effect might be expected in patients, thus affecting the biomarker quality of resting-state acquisitions for clinical purposes.

Because resting-state scans are sometimes acquired before and sometimes after other behavioral or neuroimaging tasks, we – for the first time – compare systematically pre- vs. post-task resting-state differences in clinical populations. Specifically, this study investigates how resting-state functional connectivity changes in smokers and spider-fearful individuals from before to after a cue-reactivity exposure task that consisted of presenting nicotine or spider cues, respectively. In patient populations, cue-reactivity tasks are frequently used in neuroimaging paradigms to identify physiological and neural correlates of craving or anxiety, respectively. The presentation of relevant cues is also key to exposure-based interventions, which is one of the most prominent therapeutic approaches in psychiatry. Being exposed to such cues changes the psychological state of the participant, as it will increase craving in smokers or affect the anxiety levels in phobic populations. This change will likely manifest in subsequent resting-state measures. We first tested the hypothesis that resting-state connectivity is altered in clinical populations by behavioral tasks in smokers. To test whether our results were specific to smokers or whether they generalize to other clinical populations, we applied the same analysis to an independent dataset of spider fearful individuals.

## 2. Material and methods

### 2.1. Participants

#### Nicotine use dataset

We recruited 32 participants with DSM-5 criteria for nicotine use disorder (age: 26.0±5.3; gender: 16F, 15M, 1 non-binary; sex: 17F, 15M; daily cigarette consumption: 11.5±5.6, smoking history: 7.4±4.8 years of smoking, Fagerström Test for Nicotine Dependence (FTND) score = 2.8±1.8). We instructed participants to abstain from smoking for one hour before the experiment. Exclusion criteria were the use of non-cigarette tobacco substitutes such as nicotine patches, mental or neurological disorders, and MRI-incompatibility criteria (metal implants, pregnancy, etc.). The study was conducted at the Psychiatric University Hospital of Zurich and was approved by the local ethics committee of the Canton of Zurich. Three participants were excluded due to high scanner motion, resulting in a final analysis sample of N = 29.

#### Spider phobia dataset

This dataset is a subset of a larger study, of which the MRI acquisition was still ongoing at the time we performed this analysis (N = 38 as of June 15^th^, 2022). Participants were individuals with sub-clinical spider phobia, which we defined as having a Spider Anxiety Screening (SAS, Rinck et al., 2002) score above or equal to 8 (age: 22.4 ± 3.8; gender: 30F, 7M; sex: 30F, 7M); Fear of Spider Questionnaire (Szymanski & O’Donohue, 1995) score: 51.3±21.0). Exclusion criteria were MRI-incompatibility and mental or neurological disorders. This study was conducted at the University of Vienna and was approved by the ethics committee of the University of Vienna. One participant was excluded due to motion, resulting in a final sample of N = 37 for imaging analyses.

For both studies, participants gave informed written consent and received financial compensation.

### 2.2. Experimental design

#### Nicotine use dataset

We collected two 7 minutes long resting-state runs, for which we instructed the participants to let their mind wander while looking at a fixation dot. Between the two resting-state runs, the participants underwent a smoking cue-reactivity task, which consisted in viewing 330 craving-inducing pictures from the Smoking Cue Database (Manoliu et al., 2021) and other smoking databases. In total, the pre-task and the post-task resting-state runs were separated by a 20-minute-long cue-reactivity task. We refer the reader to Haugg et al., 2022 for more details regarding the cue-reactivity task. We assessed smoking urge levels with the German version of the Questionnaire on Smoking Urges (QSU-G, Toll et al, 2006), once before the scanning session and a second time after the scanning session.

#### Spider phobia dataset

The experimental design of the spider cue-reactivity study was analogous to the nicotine use dataset described previously. Spider phobics were passively exposed to 300 pictures of spiders or neutral pictures, divided into 5 runs of about 7 minutes. 6 button-press catch trials were also randomly included in each run to increase the attention and the engagement of the participant. After the first and the last passive-viewing runs, we asked the participants how tired they felt on a scale of 1 to 10. However, due to technical issues with the interphone (e.g., drop in sound quality), we collected pre-post tiredness data for N = 26 participants only. Before and after the passive-viewing task, we acquired resting-runs of about 9 minutes each for which we asked the participants to relax and look at the fixation cross. Approximately 35 minutes elapsed between the end of first rest period and the start of the second one. Of note, all the participants in this group performed a Behavioral Avoidance Test (BAT) prior to being scanned, followed by a 15-minute walk to the scanning site.

### 2.3. MRI acquisition parameters

#### Nicotine use dataset

MR scans were collected with a 3T Philips Achieva system (Philips Healthcare, Best, The Netherlands) using a 32 phased-array head coil at the Psychiatric University Hospital, Zurich. The two resting-state functional scans were acquired with a T2*-weighted gradient-echo planar imaging (EPI) sequence (repetition time (TR) = 2000ms, echo time (TE) = 35ms, flip angle (FA) = 82°, 33 slices, no slice gap, voxel size = 3 × 3 × 3 mm^3^, field of view (FoV) = 240 × 240 × 99 mm^3^, total scan duration = 7:12 min per run). A high-resolution anatomical T1-weighted scan was acquired (FA = 8°, 237 slices, voxel size = 0.76 × 0.76 × 0.76 mm^3^, FoV = 255 × 255 × 180 mm^3^) at the end of the session.

#### Spider phobia dataset

The scans were acquired using a 3T Siemens Magnetom Skyra (Siemens, Erlangen, Germany) with a 32 □ channel head coil at the University of Vienna. Resting-state scans were collected using a multiband accelerated T2*□ weighted echo planar imaging (EPI) sequence (56 slices, no slice gap, multiband acceleration factor = 4, TR = 1250 ms, TE = 36 ms, FA = 65°, FOV = 192 × 192 × 146 mm^3^, voxel size = 2 × 2 × 2.6 mm^3^, total scan duration = 08:51 min). Structural images were acquired with a magnetization□ prepared rapid gradient □ echo (MPRAGE) sequence (FA = 8°, 208 slices, voxel size = 0.8 × 0.8 × 0.8 mm, FOV = 263 × 350 × 350 mm^3^) To decrease head motion, we taped the forehead of the participant to each side of the MRI head coil, which works by providing tactile feedback when moving (Krause et al., 2019). Additionally, we used an eye-tracker to verify that participants were not falling asleep.

### 2.4. MRI preprocessing

#### Nicotine use dataset

All analyses were performed using the CONN20b toolbox (Whitfield-Gabrieli & Nieto-Castanon, 2012) which ran on MATLAB2018a (The MathWorks Inc, Natick, Massachusetts, USA) and SPM12 (Wellcome Trust Centre for Neuroimaging, London, United Kingdom). Resting-state data was preprocessed using CONN’s default preprocessing pipeline (labeled as “default preprocessing pipeline for volume-based analyses (direct normalization to MNI-space)) and included functional realignment and unwarp, slice-timing correction, ART-outlier identification, functional and anatomical normalization into standard MNI space and segmentation into grey matter, white matter, and CSF. Functional smoothing was performed with a 6 mm FWHM Gaussian kernel. Three participants with high motion (defined as mean framewise displacement (Power et al., 2012) above 0.3 mm and/or less than 5 minutes of valid scans for each resting-state run after outlier censoring) were excluded from further analyses. This resulted in N = 29 participants.

#### Spider phobia dataset

This dataset was preprocessed using fMRIPrep 20.2.6 (Esteban et al., 2019), which is based on Nipype 1.7.0 (Gorgolewski et al., 2011).

Anatomical T1-weighted (T1w) images were corrected for intensity non-uniformity (INU) and skull-stripped. Anatomical segmentation of cerebrospinal fluid (CSF), white-matter (WM) and gray-matter (GM) was performed on the brain-extracted T1w using Fast (FSL 5.0.9, Zhang et al., 2001), and spatial normalization to MNI152NLin6Asym was performed through nonlinear registration with antsRegistration (ANTs 2.3.3), using brain-extracted versions of both T1w reference and the T1w template.

Functional preprocessing was performed as follows: generation of a reference volume and its skull-stripped version using a custom methodology of fMRIPrep, distortion correction of the reference based on a phase-difference fieldmap, co-registration of the reference volume to the subject’s T1w space using FLIRT (FSL 5.0.9, Jenkinson & Smith, 2001), configured with nine degrees of freedom. Next, head-motion parameters with respect to the reference volume (transformation matrices, and six corresponding rotation and translation parameters) were estimated using MCFLIRT (FSL 5.0.9, Jenkinson, 2002), after which the BOLD time-series were slice-time corrected using AFNI (Cox, 1996). The corrected BOLD time-series were then resampled to MNINLin6Asym space as a single interpolation step that combines transformation parameters that were previously estimated (motion corrected transformation, field distortion corrected warp, BOLD-to-T1w transformation, and T1w-to-MNI template transformation) using antsApplyTransforms (ANTs 2.3.3), configured with Lanczos interpolation.

Framewise Displacement and anatomical component-based noise regressors (aCompCor, Behzadi et al., 2007, performed with fMRIprep’s custom thresholds) were also estimated from CSF and the WM segmentation maps.

An additional spatial smoothing step with a Gaussian kernel of 6 mm FWHM was performed in CONN. One participant with high motion (mean FD > 0.3 mm) was excluded, resulting in a total number of 37 participants for further analysis.

### 2.5. Denoising

While preprocessing pipelines differed, the time courses were denoised using the same pipeline in CONN. Preprocessed data was denoised with a linear regression method, which included an aCompCor noise correction procedure with the first five principal components of WM masks and CSF masks as noise regressors as well as realignment parameters and ART-derived outliers regressors. Following CONN’s standard denoising pipeline, we applied default band-pass filtering (0.008-0.09 Hz) and linear de-trending.

### 2.6. ROI-to-ROI functional connectivity analysis

#### ROI mask description

To test for pre-post resting-state changes, we performed an ROI-to-ROI functional connectivity analysis of brain regions involved in substance-use disorders (Koob & Volkow, 2016) as ROIs for the smokers dataset. We used the same ROIs for the phobia dataset because our primary intention was to check for the generalizability of the findings, and also because many of these regions are also associated with anxiety disorders. Specifically, we included anatomical masks of the anterior cingulate cortex (ACC), medial prefrontal cortex (mPFC), bilateral amygdala, caudate, hippocampus, insula, nucleus accumbens, and thalamus (8 ROIs in total). All the masks were provided by the CONN toolbox and corresponded to regions of the Harvard-Oxford atlas, excepted for the mPFC, which was derived from ICA analyses based on Human Connectome Project data (Van Essen et al., 2013) (Figure 1) because we found no equivalent in the Harvard-Oxford atlas.

**Figure 1.**
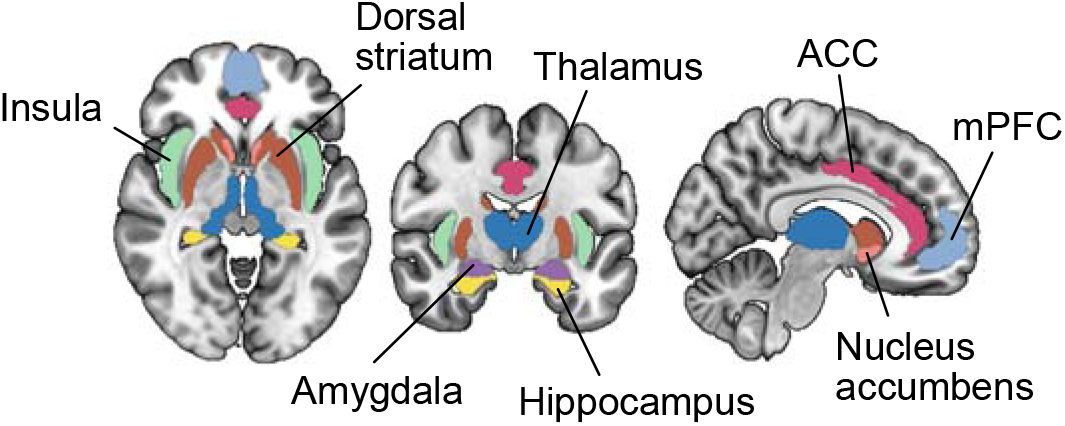
Brain regions used as seeds in seed-based analyses and ROI-to-ROI analyses. Masks for the bilateral amygdala, dorsal striatum (putamen and caudate), ventral striatum/nucleus accumbens, thalamus, insula, hippocampus and anterior cingulate cortex (ACC) were taken from the Harvard-Oxford atlas. A mask for the medial prefrontal cortex (mPFC) was derived from the Brain Connectome Project dataset and provided by the CONN toolbox.

#### First-level analysis

The signal was averaged across voxels within a mask, resulting in one time course per ROI. Pearson’s r correlation values between time courses were then computed for each pair of ROIs, and Fischer-transformed. We will refer to them as resting-state functional connectivity (rsFC) values.

#### Second-level analysis

We performed two-tailed paired t-tests comparing pre-task and post-task rsFC maps and applied Bonferroni correction to control for multiple testing (28 tests).

### 2.7. Seed-based functional connectivity analysis

In addition to ROI-to-ROI analyses, we performed seed-based analyses with each ROI as an independent seed. Seed-based analysis differs from ROI-to-ROI as it allows probing for rsFC between a seed and brain clusters outside of our selected brain regions without prior spatial averaging over voxels inside predefined masks. Here, we averaged the time courses within a seed, and we computed the correlation value between the seed and each voxel of the whole brain. We used paired t-tests comparing pre-task and post-task rsFC maps, and used CONN’s default significance threshold criterion (uncorrected voxel-wise p < 0.001). To select clusters for which rsFC differs significantly at post-task compared to pre-task, we chose CONN’s default False Discovery Rate (FDR) correction with an additional Bonferroni correction at the cluster level to correct for multiple seeds testing (cluster p-FDR-corrected < 0.05/8 = 0.00625).

### 2.8. Post hoc comparison between within-subject resting-state changes vs. resting-state differences between clinical groups

The aim of this analysis serves to assess the magnitude of task-induced pre-post resting-state changes within a clinical group in comparison to resting-state differences between the clinical groups (spider phobia vs. smokers). For this, we selected the connectivity values that displayed significant pre-post changes during the ROI-to-ROI analysis and fit a linear mixed-effects model with phobia vs. smokers group (clinical trait) as a fixed effect, and pre vs. post (psychological state) as both a fixed effect and repeated effect. The analysis was performed using the MIXED procedure in IBM SPSS Statistics, version 28 (IBM Corp., Armonk, N.Y., USA).

## 3. Results

### 3.1. Behavioral changes

#### Nicotine use dataset

Craving levels as assessed by the QSU right before scanning (pre-urge = 10 ± 31) and right after scanning (post-urge = 47 ± 30) increased considerably (p < 0.0001, Wilcoxon’s signed rank test).

#### Spider phobia dataset

Tiredness, as assessed orally after the first (pre-tiredness = 3.1 ± 2.0) and after the last run of the passive-viewing task (post-tiredness = 5.7 ± 1.6), also increased considerably (p < 0.0001, Wilcoxon’s signed rank test).

### 3.2. ROI-to-ROI analysis results

In the nicotine use dataset, the ROI-to-ROI analysis showed a pre-post decrease of rsFC between the following pairs of ROIs (Figure 2A): thalamus - insula (p-unc = 0.0040), thalamus - amygdala (p-unc = 0.0206), as well as nucleus accumbens - mPFC (p-unc = 0.0285), and an increase of rsFC between: hippocampus - amygdala (p-unc = 0.0073), and hippocampus - insula (p-unc = 0.0112) (Figure 2A). However, no detected differences survived Bonferroni correction (p > 0.05/28 = 0.0018).

**Figure 2.**
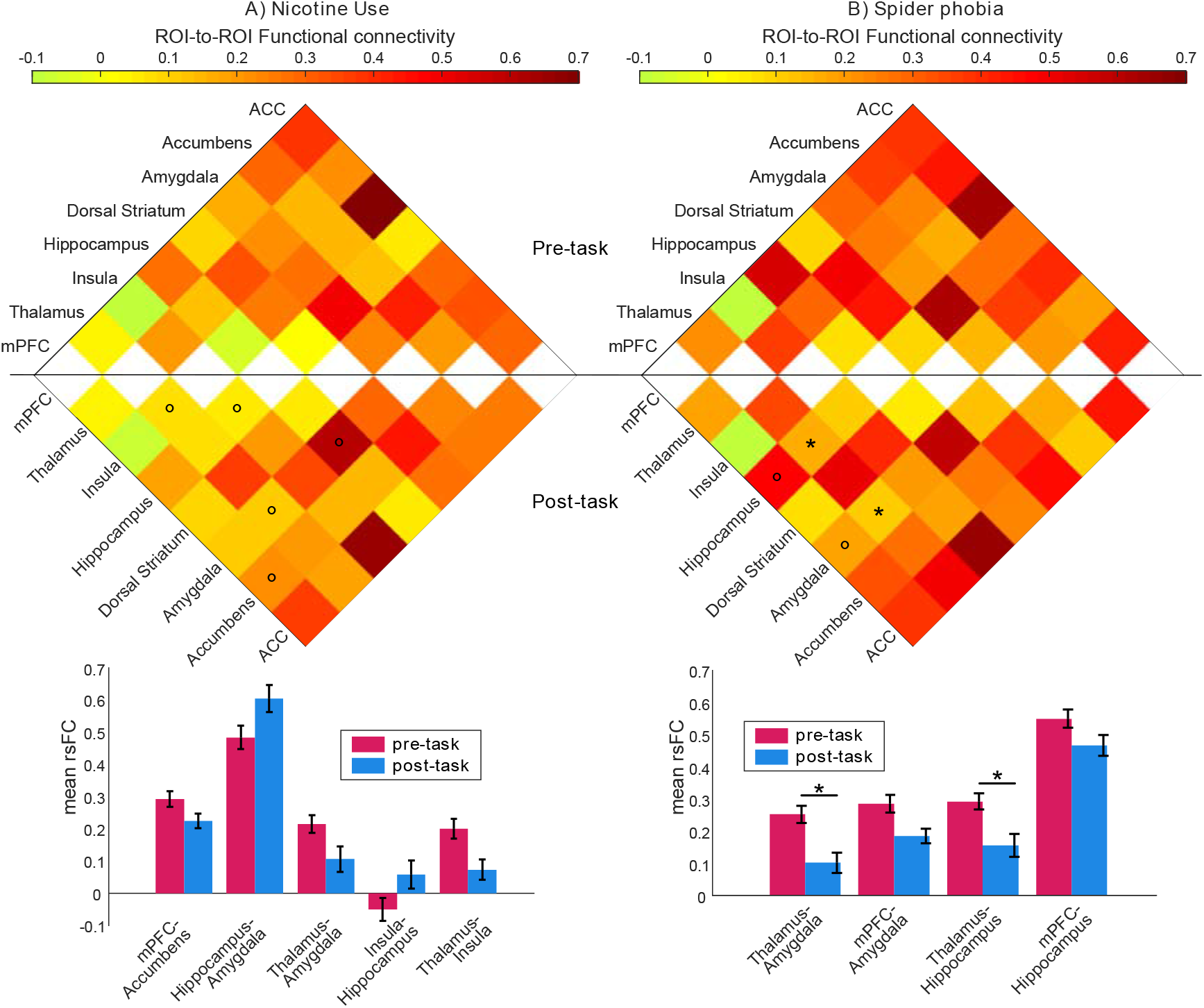
ROI-to-ROI pre-post functional connectivity matrix and bar plots. The upper part of the matrix contains rsFC before the task, and post-task rsFC is mirrored on the lower half. (A) In the nicotine use disorder dataset, paired-t-tests (°p-unc < 0.05, two-tailed) comparing pre and post task rsFC showed a decrease in nucleus accumbens–mPFC, thalamus-amygdala, thalamus-insula, and an increase in hippocampus-amygdala and insula-hippocampus, but none survived Bonferroni correction. (B) In the spider phobia dataset, mPFC-amygdala, mPFC-hippocampus, thalamus-amygdala, and thalamus-hippocampus rsFC significantly decreased, with the last two connections being significant after Bonferroni correction (*p < 0.05/28 = 0.0018). Abbreviations: medial prefrontal cortex (mPFC), anterior cingulate cortex (ACC). Error bars in bar plots correspond to SEM. N_smokers_ 29. N_phobia_ 37.

In the spider phobia dataset, we found a significant decrease of thalamus - amygdala rsFC (p-unc = 0.0003, p-corr = 0.009) and thalamus – hippocampus rsFC (p-unc = 0.0016, p-corr = 0.0448). There was also a decrease of amygdala - mPFC rsFC (p-unc = 0.0053) and hippocampus - mPFC rsFC (p-unc = 0.0480) but these two did not survive Bonferroni correction (Figure 2B).

### 3.3. Seed-based analysis results

In the nicotine use dataset, pre-post task rsFC changes comprised (1) decreased rsFC between the insular seed to a cluster in the right thalamus, (2) decreased rsFC between the dorsal striatal seed and a cluster in the lingual gyrus, and (3) increased rsFC between the mPFC seed to clusters in various cortical areas (e.g supramarginal gyrus, precentral gyrus) and a decreased rsFC to clusters in the cerebellum.

In the spider phobia dataset, we found (1) decreased rsFC between the ACC seed and cortical regions, and increased rsFC with a cerebellar cluster, (2) decreased rsFC between the amygdala seed and a thalamic cluster, (3) increased rsFC between the dorsal striatal seed to clusters in the cerebellum and decreased connectivity to several cortical areas, and (4) increased rsFC between the hippocampus and cortical clusters including visual areas.

Finally, in both datasets, we found a large significant decrease in connectivity between the thalamic seed and clusters in cortical regions, including the middle temporal gyrus, supplementary motor area, anterior cingulate cortex, precuneus and insula. In addition, the spider phobia dataset showed significantly increased thalamo-cerebellar connectivity and intra-thalamic connectivity during the post run as compared to the pre run. We also found increased intra-thalamic connectivity in the nicotine use dataset (albeit uncorrected for multiple seeds testing). All clusters are reported in Table 1 and Table 2 and illustrated in Figure 3.

**Table 1.**
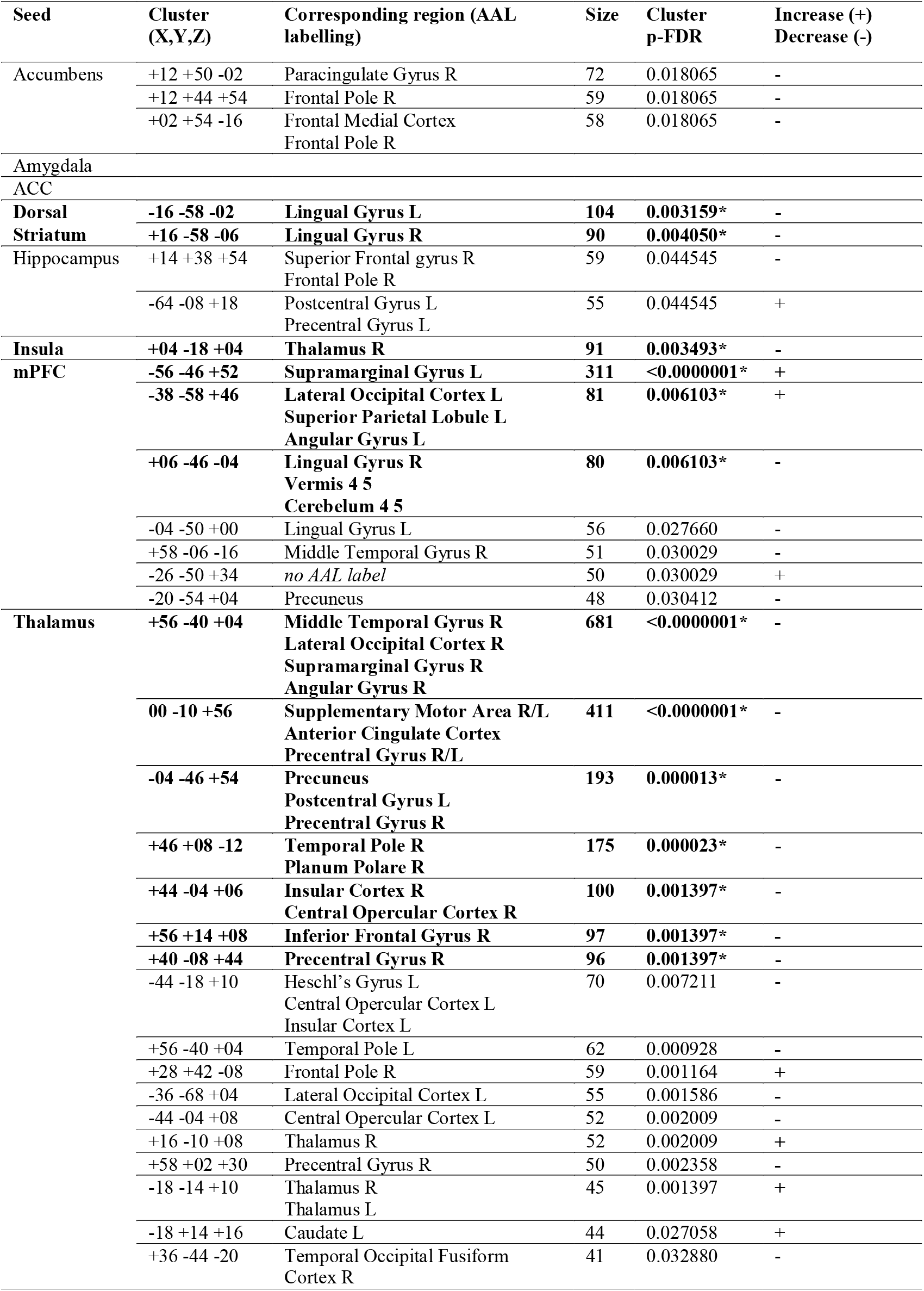
Seed-based functional connectivity changes following a smoking cue-reactivity task. Clusters were labelled using the Automated Anatomical Labeling (AAL) Atlas by the CONN toolbox. The table reports all clusters with p-uncorrected < 0.001 at the voxel level and p-FDR < 0.05 at the cluster level. *clusters that survive an additional Bonferroni correction to control for multiple seeds testing at the cluster level (p-FDR < 0.05/8).

**Table 2.**
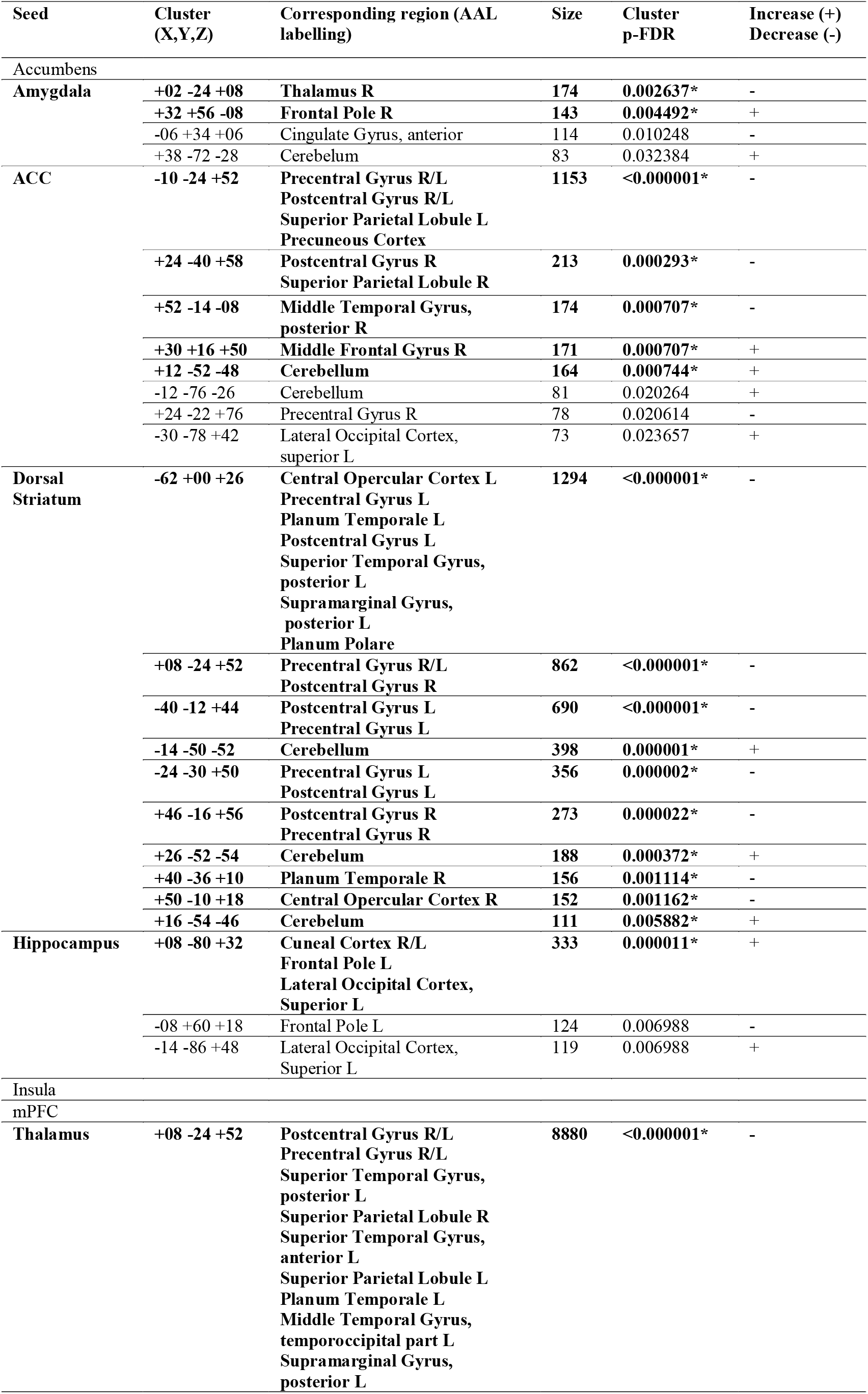
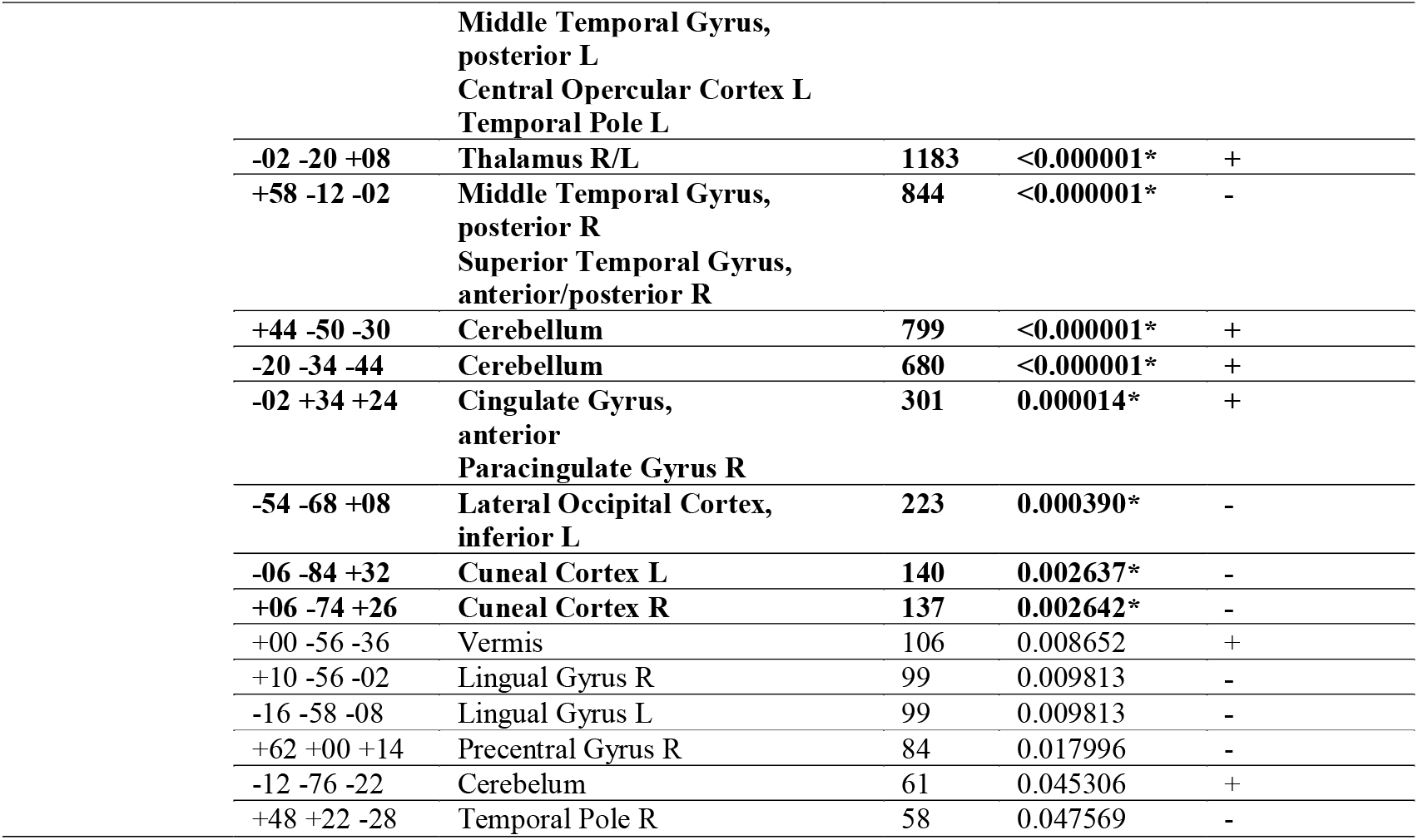
Seed-based functional connectivity changes following exposure to spider stimuli. Clusters were labelled using the Automated Anatomical Labeling (AAL) Atlas by the CONN toolbox. The table reports all clusters with p-uncorrected < 0.001 at the voxel level and p-FDR < 0.05 at the cluster level. *clusters that survive an additional Bonferroni correction (p-FDR < 0.05/8) to control for multiple seeds testing at the cluster level.

**Figure 3.**
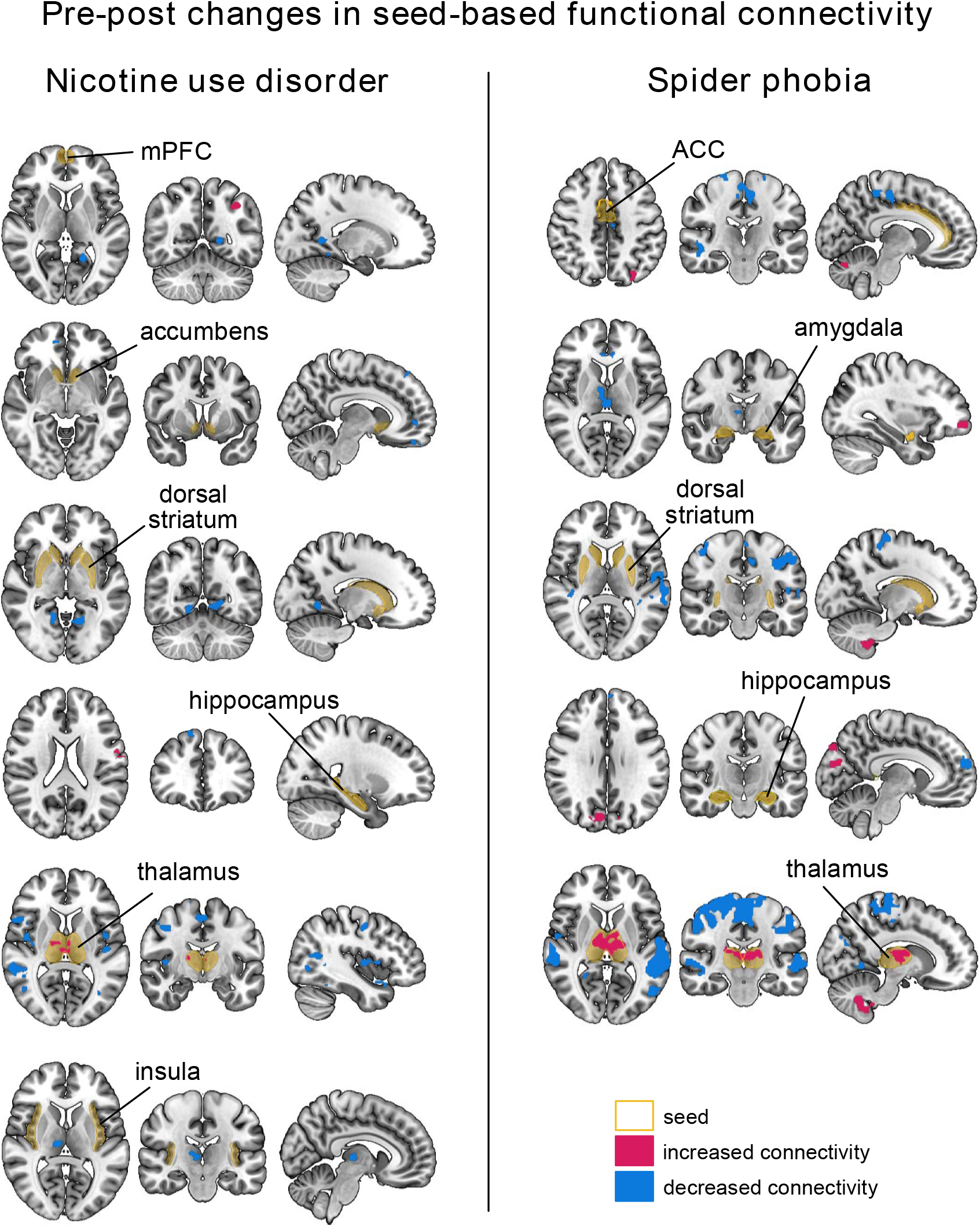
Seed-based functional connectivity changes following a smoking cue-reactivity task and a phobic cue-reactivity task in nicotine use disorder and spider phobia, respectively. Voxel-level p < 0.001 and cluster-level p-FDR < 0.05 without correction for multiple seeds were used here as significance threshold for illustrative purposes. Clusters that survived an additional Bonferroni correction for multiple seed testing are reported in Table 1 and 2.

### 3.4. Post hoc comparison between within-subject resting-state changes vs. resting-state differences between clinical groups

In the ROI-to-ROI analysis, we found significant pre-post changes in thalamus-amygdala rsFC (p-unc = 0.0003; p-corr = 0.009) and in thalamus-hippocampus rsFC (p-unc = 0.0016, p-corr = 0.0448) in the spider phobia group using paired-t-tests. For the thalamus-hippocampus connection, a linear mixed-effects model revealed a significant main effect of both clinical group (F = 13.6; p < 0.001) and state (F = 11.6; p = 0.001). For the thalamus-amygdala connection, however, there was a significant main effect of the psychological state (i.e. pre vs. post task) (F = 20.1; p < 0.001) but no significant effect of the clinical trait (i.e. smokers vs. phobics) (F = 0.451; p = 0.504), indicating that pre-post changes were even greater than differences between groups for this pair.

**Figure 4.**
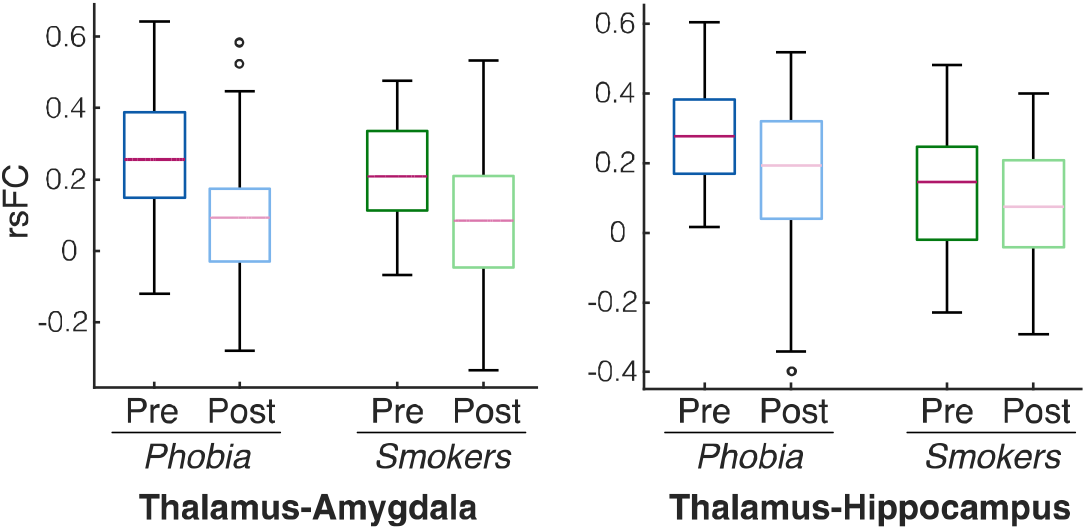
Comparison between within-subject changes (pre vs. post task) and between-group differences (phobics vs. smokers). Pre-post change of thalamus-amygdala rsFC exceeded the difference between groups.

## 4. Discussion

Previous studies have shown that behavioral tasks (Cecchetto et al., 2019; Sarabi et al., 2018) can affect fMRI resting-state functional connectivity of healthy populations, but such alterations have never been investigated in clinical populations. Here, we compared resting-state functional connectivity before and after a smoking cue-reactivity task in smokers and a spider cue-reactivity task in spider phobics. We found significant rsFC alterations in the nicotine use disorder dataset when the mPFC, insula, dorsal striatum and thalamus were defined as seeds, and in the spider phobia dataset, when the amygdala, ACC, dorsal striatum, hippocampus, and thalamus were defined as seeds. Of note, in both datasets, we found a decreased rsFC between the thalamic seed and cortical areas, as well as increased rsFC with a cluster within the thalamus itself, which indirectly reflects an increase in thalamic regional homogeneity. Finally, thalamus-amygdala and thalamus-hippocampus ROI-to-ROI rsFC were significantly reduced for spider phobics, and this thalamus-amygdala rsFC reduction in spider phobics was even greater than the difference with rsFC of smokers.

Resting-state data has long been a potential candidate for identifying clinical biomarkers of mental disorders, including tobacco use disorder (John R. Fedota & Stein, 2015), with the hope that tracking treatment outcome and stratifying disorders into subtypes would help designing personalized treatment plans. However, the field suffers from important drawbacks: uncertainty of machine-learning target labelling (e.g. psychiatrists can disagree when assigning a diagnosis to a patient), unclear boundaries between psychiatric disorders, mismatch between disorder definition, symptoms and neural underpinnings, etc. (Parkes et al., 2020; Yamada et al., 2017). Further, single-study findings tend to poorly generalize across multiple studies, an issue that has been partly attributed to site-specific technical characteristics (imaging sequence, scanner type, field strength…) or differences in data acquisition protocols (Yamashita et al., 2019). Besides these technical aspects, differences in psychological states can also contribute to data heterogeneity. Our results in two different clinical or subclinical populations corroborate this view, as task-induced increase of craving and phobic alertness were accompanied with large resting-state changes in disorder-relevant brain regions. In particular, for the thalamus-amygdala rsFC, within-subject pre-post alterations even exceeded the difference between clinical groups during pre-rest periods. This is remarkable since, between these two groups, there were major differences in scanner type, acquisition parameters, and preprocessing pipeline. This indicates that this connection, known for being highly relevant for addiction (Rich et al., 2019) and fear processes (Silva et al., 2021), may be more sensitive to psychological states than to clinical traits.

In the spider phobia group, other alterations include a decreased hippocampus-thalamus ROI-to-ROI rsFC, as well as an increase of hippocampal seed connectivity with visual areas, which could reflect hippocampal reorganization related to stress, fear retrieval or fear extinction due to being exposed to aversive stimuli (Chang & Yu, 2019). The dorsal striatum also becomes connected to somatosensory cortical areas (postcentral gyrus), and motor control areas (precentral gyrus), which might be linked to fight-or-flight mechanisms or inhibitory control mechanisms following fear exposure (Stanley et al., 2021). In the smokers group, other seed-based connectivity alterations include a decreased connectivity between the thalamus and regions such as the precunous, the ACC and the insula. This is in line with previous studies that contrasted smokers and non-smokers (Chaoyan Wang et al., 2018), or relapsers vs. non-relapsers (Chao Wang et al., 2020), which indicates that these brain changes might be related to psychological changes in the nicotine use disorder patients (e.g. increased craving).

However, the pre-post changes of psychological and neural states are not uniquely driven by the clinical specificities of the task, and the change of connectivity does not exclusively have to be attributed to changes of urge to smoke or fear states. Many other psychological states vary from pre to post periods, among which cognitive fatigue, tiredness, or sleepiness at the end of a scanning session, as well as hyper-vigilance and hyper-attention to sensory bottom-up information at the beginning of the session are not uncommon. This is notably illustrated by an increase of self-reported tiredness scores over the course of scanning. The similarity between changes in thalamic-based connectivity in both datasets, namely increase of intra-thalamic connectivity and decrease of cortico-thalamic connectivity, is quite remarkable. Among many other processes, the thalamus is known for playing a key role in regulating sleep-wake cycles (Scammell et al., 2017). Akin to our study, human (Hale et al., 2016) and animal studies (Sysoev et al., 2021) also found both decreased thalamo-cortical and increased intra-thalamic connection when tracking participants or mice during the process of falling asleep. Further, decreased thalamo-cortical connectivity has been associated with unconsciousness induced by anesthesia (Akeju et al., 2014) and being sleep-deprived (Shao et al., 2013), whereas symmetrically, increased thalamo-cortical functional connectivity has been linked to chronic insomnia (Kim et al., 2021). Considering the consistency of the results across a wide range of consciousness/sleep-related operationalizations, this decrease of thalamo-cortical connectivity has been proposed as a solid hallmark for changes of consciousness states (Picchioni et al., 2014). This indicates that our thalamic connectivity change likely reflects a reduction of vigilance (using an eye tracker, we did not see any of the participants of the spider phobia study fall asleep) which may further confound biomarker detection. However, the current data is not conclusive if the vigilance reduction is due to patients having been in a hyper-vigilant state during pre-task resting-state scans, due to vigilance reducing below baseline levels during post-task resting-state scans, or both.

### Limitations

First, the clinical interpretability of our disorder-specific findings is limited by the lack of additional control groups. For instance, to find rsFC changes that are specific to the smoking task or the smoking population, a control group of smokers exposed to non-smoking related pictures would be required. Second, to firmly establish the interpretation of the thalamus-related changes that we observed in both groups are associated with changes in vigilance, more rigorous measures on vigilance and tiredness would be beneficial. Third, the comparability between the spider phobia group and the nicotine use group also needs to take into account the technical differences in fMRI acquisition (e.g. MR-scanner, imaging sequence), paradigm (e.g. duration of the resting-state scans, eyes closed or open), and data preprocessing (e.g. SPM12 vs. AFNI). On the other hand, the finding that the pre- vs. post-task resting-state changes are more pronounced than the resting-state connectivity differences between the two clinical groups is even more remarkable given the data acquisition, paradigm, and data analysis differences. Finally, our study included two clinical populations to generalize beyond substance use disorder. While it is likely that similar effects that are specific to the clinical condition and more general effects (such as vigilance changes) will also be found in other clinical conditions, this will need to be demonstrated in further studies.

### Conclusion

In all, this study confirms that resting-state measures in clinical populations can be substantially altered by task-induced psychological states. Hence, pooling pre-task and post-task resting-state scans for biomarker detection of stable clinical traits should consider the psychological state of the patients. This implies that when publishing and making resting-state data publicly available (Tanaka et al., 2021; Van Essen et al., 2013), the complete experimental design should be reported as standard practice. This includes a detailed description of the resting-state acquisition, and whether or not other tasks were performed before the resting-state acquisition, no matter if inside or outside the MR scanner. Even though task-free resting-state acquisitions are very suitable for pooling of data in search for potential biomarkers in psychiatry, data aggregation and interpretation of results needs to consider not just technical differences but also the psychological states of the patients.

